# Secreted ORF8 is a pathogenic cause of severe COVID-19 and is potentially targetable with select NLRP3 inhibitors

**DOI:** 10.1101/2021.12.02.470978

**Authors:** Xiaosheng Wu, Michelle K. Manske, Gordon J. Ruan, Taylor L. Witter, Kevin E. Nowakowski, Jithma P. Abeykoon, Xinyi Tang, Yue Yu, Kimberly A. Gwin, Annie Wu, Vanessa Taupin, Vaishali Bhardwaj, Jonas Paludo, Surendra Dasari, Haidong Dong, Stephen M. Ansell, Andrew D. Badley, Matthew J. Schellenberg, Thomas E. Witzig

## Abstract

COVID-19 is a significant cause of morbidity and mortality in blood cancer patients, especially those on immunosuppressive therapy. Despite extensive research, the specific factor associated with SARS-CoV-2 infection that mediates the life-threatening inflammatory cytokine response in patients with severe COVID-19 remains unidentified. Herein we demonstrate that the virus-encoded Open Reading Frame 8 (ORF8) protein is abundantly secreted as a glycoprotein *in vitro* and in symptomatic patients with COVID-19. ORF8 specifically binds to the NOD-like receptor family pyrin domain-containing 3 (NLRP3) in CD14^+^ monocytes to induce a non-canonical inflammasomal response, and a canonical response when the second activation signal is present. Levels of ORF8 protein in the blood correlate with severity and disease-specific mortality in patients with acute SARS-CoV-2 infection. Furthermore, the ORF8-induced inflammasome response was readily inhibited by the NLRP3 inhibitor MCC950 *in vitro*. Our study identifies a dominant cause of pathogenesis, its underlying mechanism, and a potential new treatment for severe COVID-19.

**Key points:**

- Secreted glycoprotein ORF8 induces monocytic pro-inflammatory cytokines involving the activation of the NLPR3 inflammasome pathway.
- ORF8 is prognostically present in the blood of symptomatic patients with covid-19 and is targetable with NLRP3 inhibitor MCC-950.

## INTRODUCTION

COVID-19, the global pandemic caused by infection with severe acute respiratory syndrome coronavirus 2 (SARS-CoV-2), has infected more than half a billion people and caused over six million deaths worldwide.^1^ Numerous studies have shown that the production of pro-inflammatory cytokines/chemokines including IL1β, IL6, IL8, and CCL2 is responsible for life-threatening symptoms.^2,3^ It is also known that the viral load, cytokine levels, and disease severity are tightly associated^4–6^, and the virus-neutralizing antibodies and IL1β pathway antagonists could readily mitigate symptoms and improve clinical outcomes.^7,8^ However, the intermediate viral factor that directly causes the inflammatory cytokine responses remains unidentified. It has been demonstrated that SARS-CoV-2 infection localizes to nasal and pulmonary epithelial cells,^9,10^ while the cytokine response is more systemic. It seems irreconcilable how this cytokine response is initiated given that no live virus has been reported in the blood based on transfusion medicine studies.^11–14^ We hypothesized that an inflammatory byproduct of SARS-CoV-2 replication is released into the bloodstream resulting in a systematic cytokine response in severe COVID-19 patients.

Upon infection of human cells, the SARS-CoV-2 virus replicates its 29.9kb RNA genome and produces up to 29 possible viral proteins, including 16 non-structural proteins (NSP1-16), four structural proteins Spike (SPK), Membrane (MEM), Envelope (ENV) and Nucleocapsid (NUC) and nine accessory proteins (ORF3A, 3B, 6, 7A, 7B, 8, 9b, 9c, and 10). While only four structural proteins along with the RNA genome are assembled into new viral particles, the other viral proteins are thought to be left behind^15^ which may disrupt host cell functions.^16,17^ Herein, we demonstrate that the SARS-CoV-2 encoded ORF8 is abundantly secreted as a glycoprotein into culture supernatant *in vitro* and into the bloodstream in patients with COVID-19. Glycosylated ORF8 stimulates CD14^+^ monocytes to produce a group of pro-inflammatory cytokines including IL1β through an NLRP3-mediated inflammasome response, and the ORF8/NLRP3 axis is targetable by NLRP3 inhibitor MCC-950.

## MATERIALS AND METHODS

### Study Samples

Human serum samples were collected from Mayo Clinic patients with a documented diagnosis of COVID-19 infection and consented to COVID-19 Research Task Force Specimen Biobank. Samples for these studies were requested and approved by the Task Force Review Committee and the Mayo Clinic Institutional Review Board. Deidentified fresh leukocyte cones of healthy donors were obtained from Mayo Clinic Blood Bank. Use of mouse splenic B cells from C57/B6 mice was approved by the Institutional Animal Care and Use Committee of the Mayo Clinic.

### SARS-CoV-2 protein constructs and their expression in human cells

Lentiviral constructs expressing SARS-CoV-2 proteins originally described by Gordon, et al ^16^ were purchased from Addgene (Supplemental Table 1), and used to make stable expression cell lines in HEK293 cells by lentiviral transduction, and transient expression in HEK293F cells with HyCell TransFx media (Cytiva) method for ORF8 protein purification. During the study, we first used the conditioned media (CM) (made by dialyzing the culture supernatants of ORF8-expressing HEK293 cells) for PBMC stimulation, it was later replaced with purified glycosylated ORF8.

### Stimulation PBMCs or THP-1 and cytokine detection

Fresh PBMCs (3×10^6^) isolated from healthy donors were stimulated with either 20% (v/v) of CM-ORF8 for 72 hours, or 200ng/ml pure ORF8 for 24 hours in a total volume of 1.5ml RPMI/10% FCS based on the titration results (Supplemental Figure 6). The cytokines expression was determined using qPCR (SYBR-green method, primers are listed in Supplemental Table 3), Luminex using a custom procarta Luminex 6-plax-cytokines (IL1β, IL6, IL8, IL18, CCL2, and TNFa) detection kit (ThermoFisher) (Supplemental Fig. 5), intracellular flowcytometry, or Western blotting. For qPCR, housekeeping gene β-Actin or HPRT1 was used for normalization.

### Detection of ORF8 protein in sera of patients newly infected with SARS-CoV-2

For detection of ORF8 protein in serum samples from patients newly infected with SARS-CoV-2, typically four microliters of serum were directly mixed with 36 microliters of 2x Laemmli buffer with 5% of BME, boiled for 3 min before load to 10-20% SDS-PAGE gels, the blots were probed with 1:5000 diluted rabbit anti-ORF8 antibody (MyBioSource, Cat# MBS3014575) at 4°C overnight. The images were developed using an ECL reagent kit.

### Analysis of ORF8 expression and survival in patients with COVID-19

Overall survival was calculated from the time of COVID-19 diagnosis, using the Kaplan-Meir method on JMP 14.0 software (SAS Institute, Cary, NC).

Additional methods are described in supplemental Materials and methods.

## RESULTS

### Secretion of SARS-CoV-2 proteins from human cells

To determine the secretion potential of SARS-CoV-2 encoded proteins from human cells, we transduced HEK293 cells with one of 22 lentiviral constructs available to us each expressing a SARS-CoV-2 protein tagged Strep II (Supplemental Table 1). The secreted viral proteins in the culture supernatants were enriched and analyzed by Western blotting, and their levels were compared to their total cellular expression in whole cell lysates. As shown in Fig. 1a, most viral proteins were robustly expressed with the expected molecular sizes except for NSP13, ORF10, MEM, and NSP4 due to their size or solubility issues. While NSP2, 5, 7, 9, 10, 12, 14, 15, ORF8, and NUC proteins were all detected as secreted proteins, ORF8 was the single most robustly secreted protein (Fig. 1b). This is consistent with *in silico* analysis (Supplemental Fig. 2) showing that all secreted NSPs possess an unconventional protein secretion signal (UPS) while ORF8 carries a classical protein signal sequence (SS) and an N-link glycosylation sites (^78^<u>N</u>YTV). These results suggest that ORF8 is efficiently secreted through the classical ER/Golgi protein secretion pathway, and its glycosylation explains the up-shifting and smearing of the ORF8 protein band (Fig. 1b).

**Figure 1.**
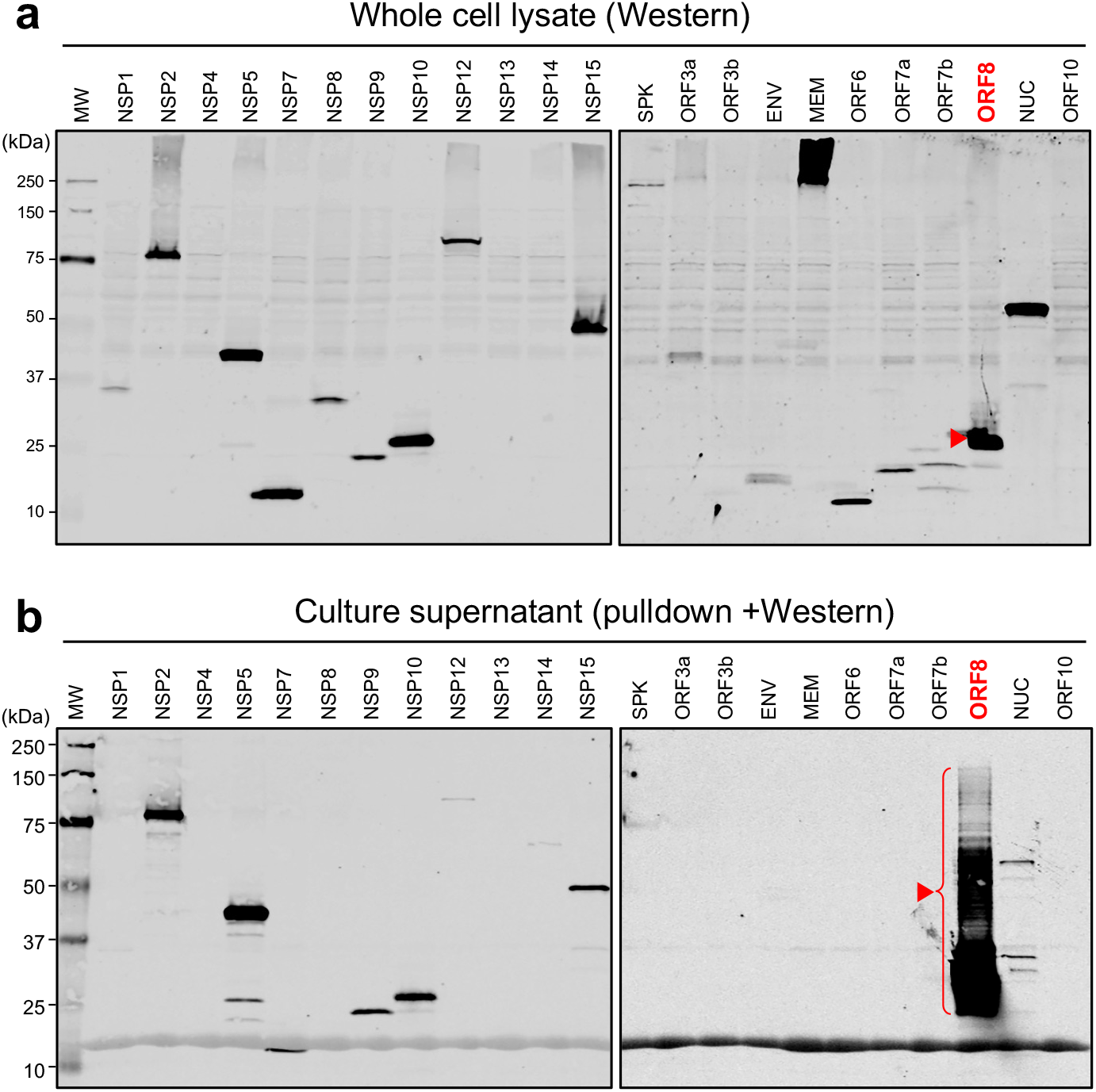
Secretion property of SARS-CoV-2 proteins. **a.** Western blot image of cell lysates for validating SARS-CoV-2 protein expression in HEK293 cells. The blots were probed with an anti-Strep II tag antibody and anti-HA (for SPK protein). **b.** Western image of SARS-CoV-2 protein pulldown using StrepTactin^™^ beads and anti-HA-beads (for SPK) from the culture supernatants for validating SARS-CoV-2 protein secretion from HEK293 cells. The blots were also probed with an anti-Strep II tag antibody and anti-HA (for SPK protein).

### Secreted ORF8 protein induces the production of pro-inflammatory cytokines

We then queried if any of the secreted SARS-CoV-2 proteins could induce the expression of pro-inflammatory cytokines seen in patients with COVID-19. We treated human peripheral blood mononuclear cells (PBMCs) from healthy donors with conditioned media (CM) containing major secreted NSPs or ORF8 proteins followed by an assessment of cytokine expression. We found that none of the secreted NSP proteins induced cytokine expression (Fig. 2a) while ORF8-containing CM (ORF8-CM) induced the expression of IL1β, IL6, IL8, and CCL2 up to 5-fold in PBMCs from select donors (Fig. 2b). Since IL1β, IL6, IL8, and CCL2 are among the key cytokines elevated in patients with severe COVID-19,^18^ it is conceivable that secreted ORF8 is the viral factor responsible for inflammatory cytokine response in patients with severe COVID-19.

**Figure 2.**
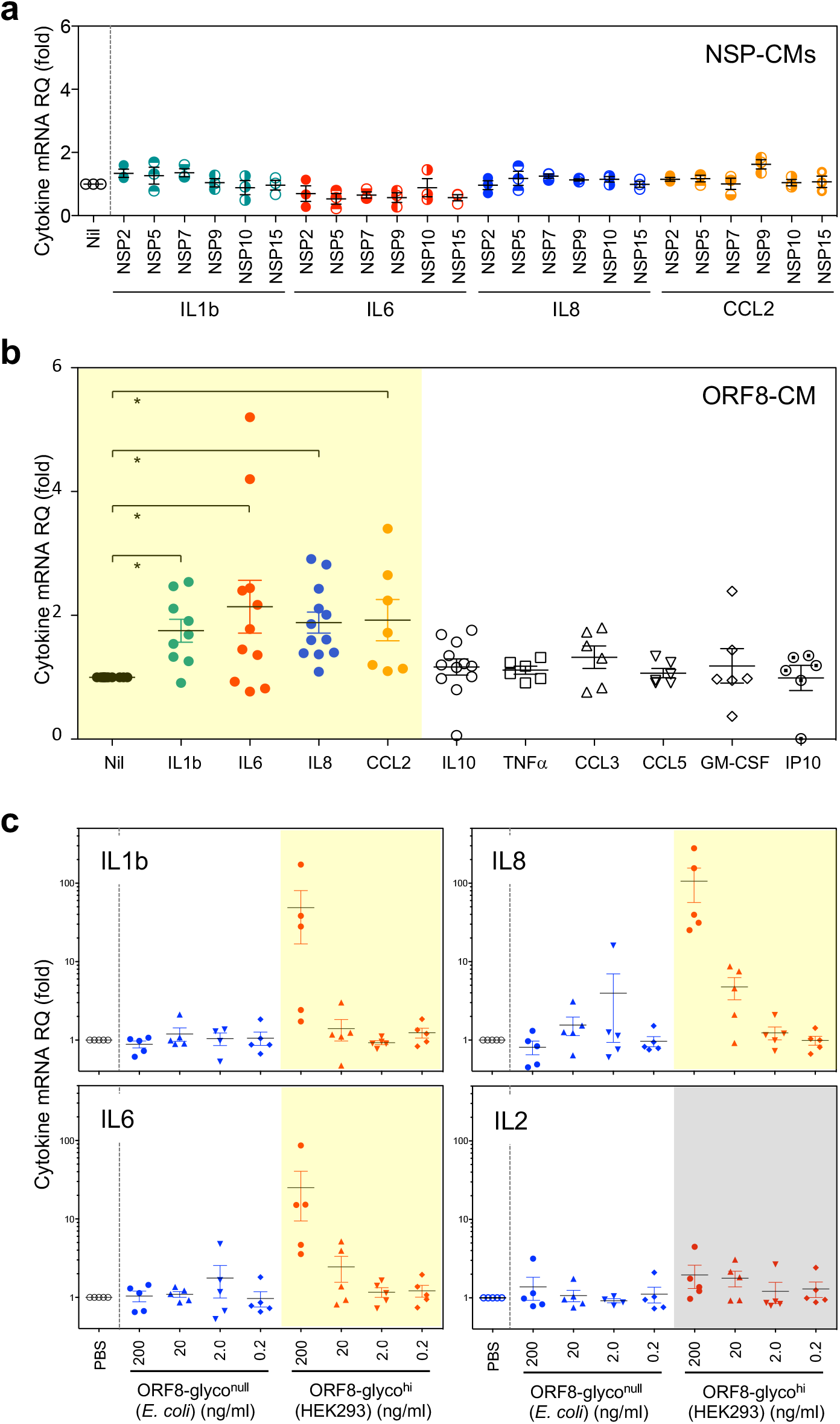
Secreted ORF8 specifically induces the expression of pro-inflammatory cytokines and is glycosylation dependent. **a.** Conditioned media containing major secreted NSP proteins did not induce pro-inflammatory cytokines in PBMCs of healthy donors; **b**. Conditioned media containing secreted ORF8 specifically induced pro-inflammatory cytokines (highlighted in yellow) IL1β, IL6, IL8, and CCL2 in PBMCs from unselected healthy donors. **c.** Purified highly glycosylated ORF8 (ORF8-glycol^hi^) from HEK293 but not unglycosylated ORF8 (ORF8-glycol^null^) from *E. coli* stimulates PBMCs to produce pro-inflammatory cytokines IL1β, IL6, IL8 but not T cell cytokine IL2.

To evaluate whether the glycosylation of ORF8 affected cytokine induction, we purified ORF8 from the HEK293 culture supernatant and from ORF8-expressing *E. coli* and designated them as ORF8-glyco^hi^ and ORF8-glyco^null^, respectively. As shown in Fig. 2c that IL1β, IL-6, and IL-8 but not IL-2 were robustly induced by the pure ORF8-glyco^hi^ in a dose-dependent manner confirming that secreted ORF8 is capable of inducing proinflammatory cytokines seen in COVID-19 patients. Interestingly, *E. coli* expressed ORF8-glyco^null^, had no cytokine induction activity even at higher doses, suggesting that proper ORF8 glycosylation is necessary for cytokine induction.

We then tested the cytokine induction activities of ORF8 from HEK293 cells treated with the Golgi inhibitor cocktail Brefeldin-A and Monensin (BFA/M) and found that while the glycosylation level was reduced (Supplemental Fig. 3) the cytokine induction activity of BFA/M-treated ORF8 was indeed altered (actually increased), suggesting that the cytokine induction activity of ORF8 is governed by the Golgi complex. Our protein localization data (Supplemental Fig 1, 2) indicate that several SARS-CoV-2 proteins are localized to Golgi membranes and may potentially modify the glycosylation and the cytokine induction activity of ORF8. Indeed, co-expressing MEM, ENV, ORF3A, or ORF7A was able to alter the molecule weight and the cytokine-inducing activities of ORF8 (Supplemental Fig 4).

### ORF8 stimulates CD14^+^ monocytes to produce pro-inflammatory cytokines/chemokines

Next, we asked which specific cell subsets in PBMCs are the primary targets of ORF8, and what is the resulting transcriptional signatures in those cells. We performed single-cell RNA sequencing (scRNA-Seq) analysis on three PBMC samples treated with ORF8-CM or control-CM. Single-cell transcriptome-based cell clustering showed that cells in cluster 5 (C5) on the t-SNE plot (Fig. 3a) were induced to express cytokines including IL1β, IL8, and CCL2 and inflammasome pathway components upon ORF8 treatment (Fig. 3b) (Supplemental Table 2). The same cytokines are known to be elevated in COVID-19 patients.^19^ The expression of CD14, CD16, CD68, and HLA-DR identifies those cytokine secreting cells as activated monocytes.^20,21^

**Figure 3.**
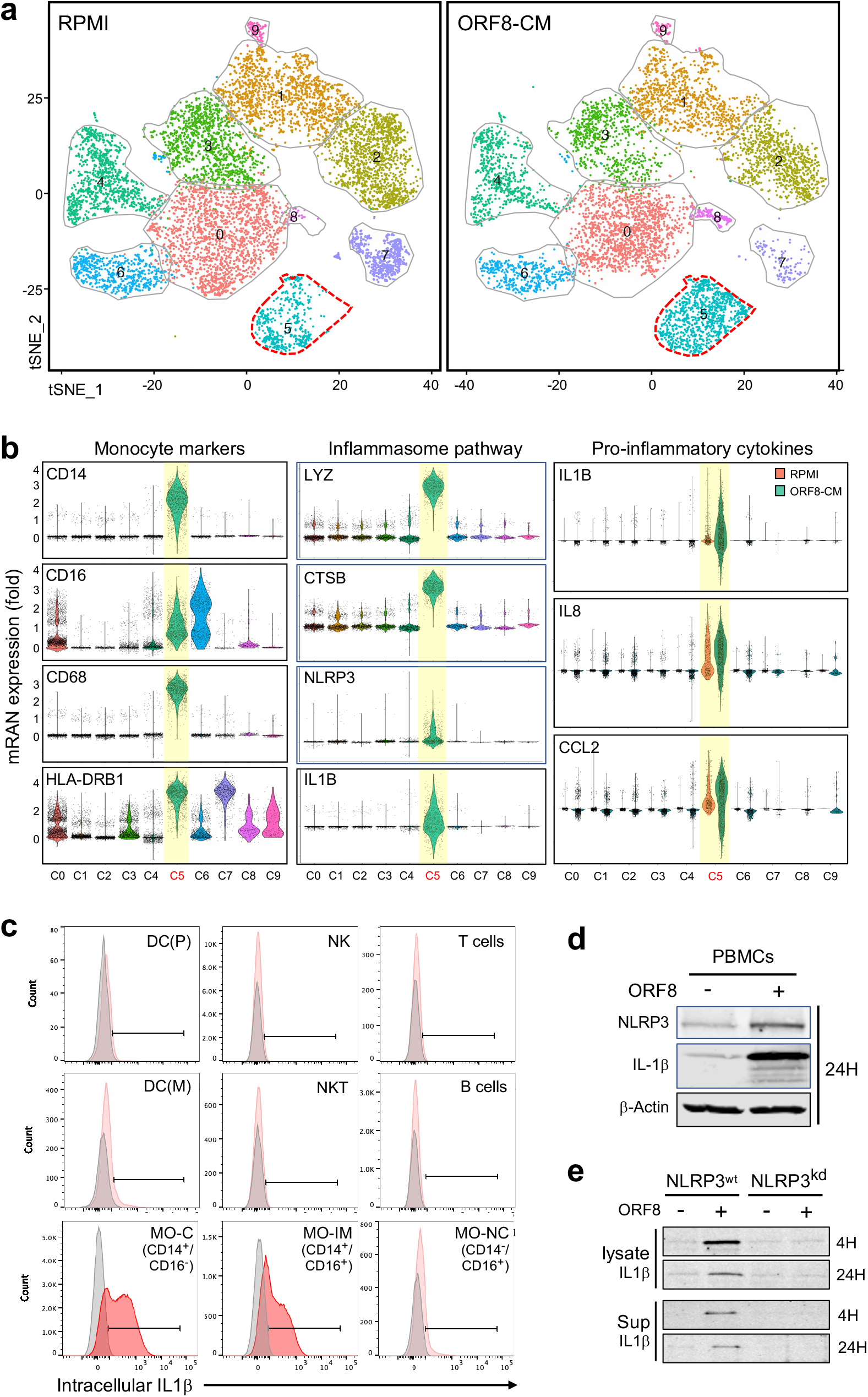
Single-cell RNA sequencing of ORF8 treated PBMCs. **a.**tSNE maps of cell clusters using concatenated scRNA-Seq data from two ORF8-responsive PBMC samples. Cells in cluster 5 were responsive to ORF8 treatment. **b.** Violin plots show the monocyte markers CD14, CD16, CD68, and HLA-DRB1, key inflammasome pathway components LYZ, CTSB, NLRP3, and IL1β, and pro-inflammatory cytokines IL1β, IL8, and CCL2 mRNA expression in cells of cluster 5. **c.** Flow cytometry analysis of intracellular IL1β protein expression in different PBMC subsets in response to ORF8 treatment. The plots show that IL1β was exclusively produced by CD14^+^CD16^-^ (classical) and CD14^+^CD16^+^ (intermediate) monocyte subsets. **d.** Western blot images validate that the induction of NLRP3 and IL1β in PBMCs upon stimulation with ORF8 indeed occurs at the protein level. **e.** Western images showing in response to ORF8 treatment, the IL1β induction in parental THP-1 cells became diminished in NLRP3 knockdown (NLRP3^kd^) THP-1 cells. The data shown are representative of at least two independent experiments except for the scRNA-Seq.

To identify which monocyte subsets are the targets of ORF8, we analyzed intracellular cytokines in ORF8 stimulated PBMCs from 15 donors by flow cytometry. Significant IL1β expression was detected in classical (CD14^+^/CD16^-^) and intermediate (CD14^+^/CD16^+^) subsets but not non-classical (CD14^low^/CD16^+^) monocytes nor cells of other lineages (B or T-lymphocytes, dendritic, NK cells) in all 15 donors (Fig. 3c). Similar results were observed for IL8 and CCL2 expression as well (Fig. 5a). Our data demonstrate that the CD14^+^ monocyte subsets are the producer of the pro-inflammatory cytokines upon ORF8 stimulation.

**Figure 4.**
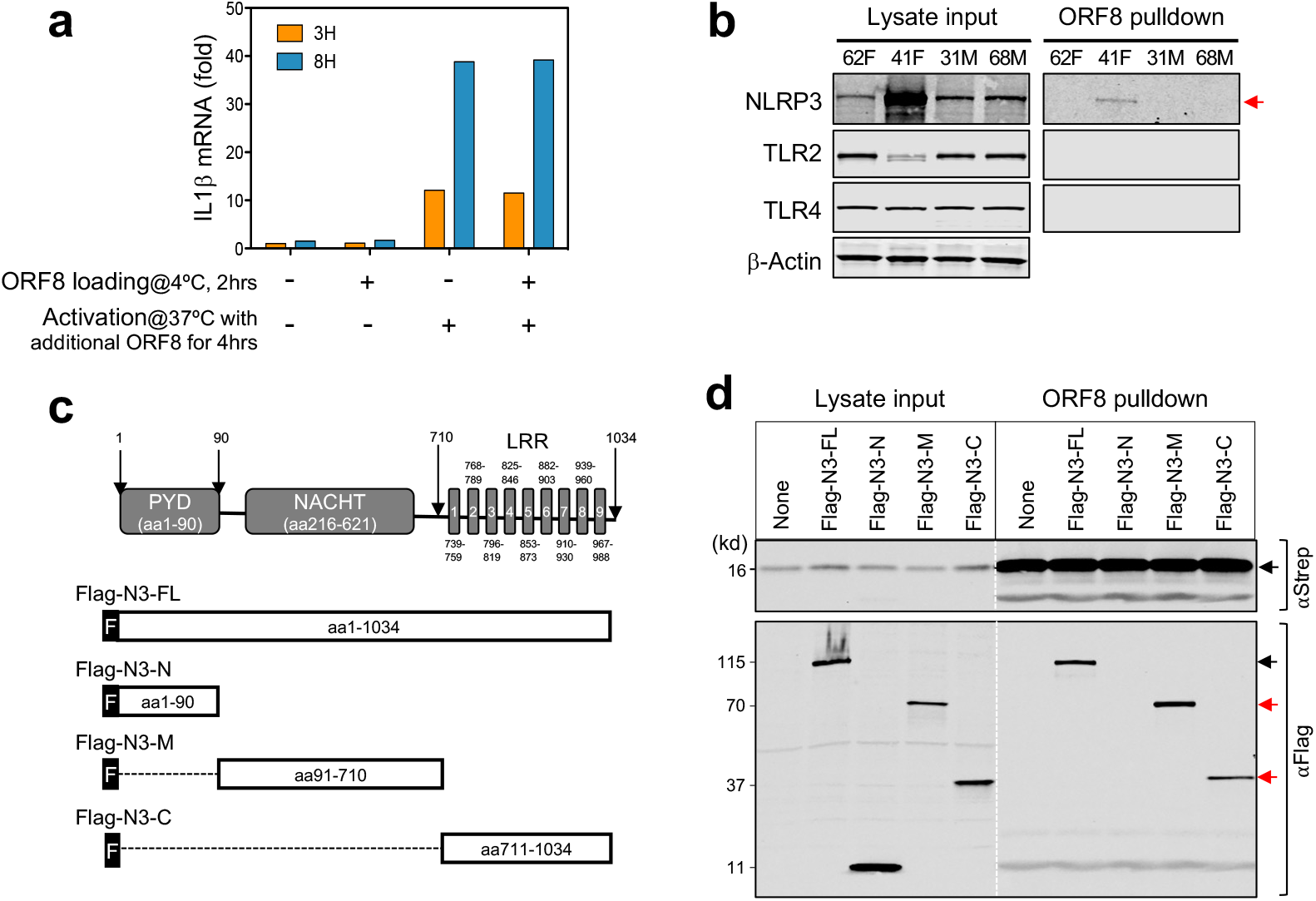
ORF8 induces proinflammatory cytokines through direct binding to NLRP3. **a.** ORF8 induces IL1β production through a non-surface-receptor-mediated process. PBMCs were pre-incubated on ice with ORF8 (500 ng/ml) for 2 hours to allow ORF8 to bind to its “surface receptors” followed by washing and incubating at 37ºC to activate the cytokine production in the presence or absence of additional ORF8 (200 ng/ml). IL1β expression was measured by qPCR. **b.** ORF8 directly binds to NLRP3 but not TLR2 or TRL4 in primary monocytes shown by affinity pulldown assay. **c.** Schematic drawing of NLRP3 deletion constructs for mapping ORF8 binding domains in NLPR3. **d.** Mapping NLRP3 domains that bind to ORF8 by affinity pulldown assay using ORF8-Strep-Tacin®beads. Western images of total cell lysates (left half) were used as expression controls and Western images of precipitated proteins (right half) showing ORF8 efficiently binds the full-length as well as two NLRP3 deletion mutant proteins.

**Figure 5.**
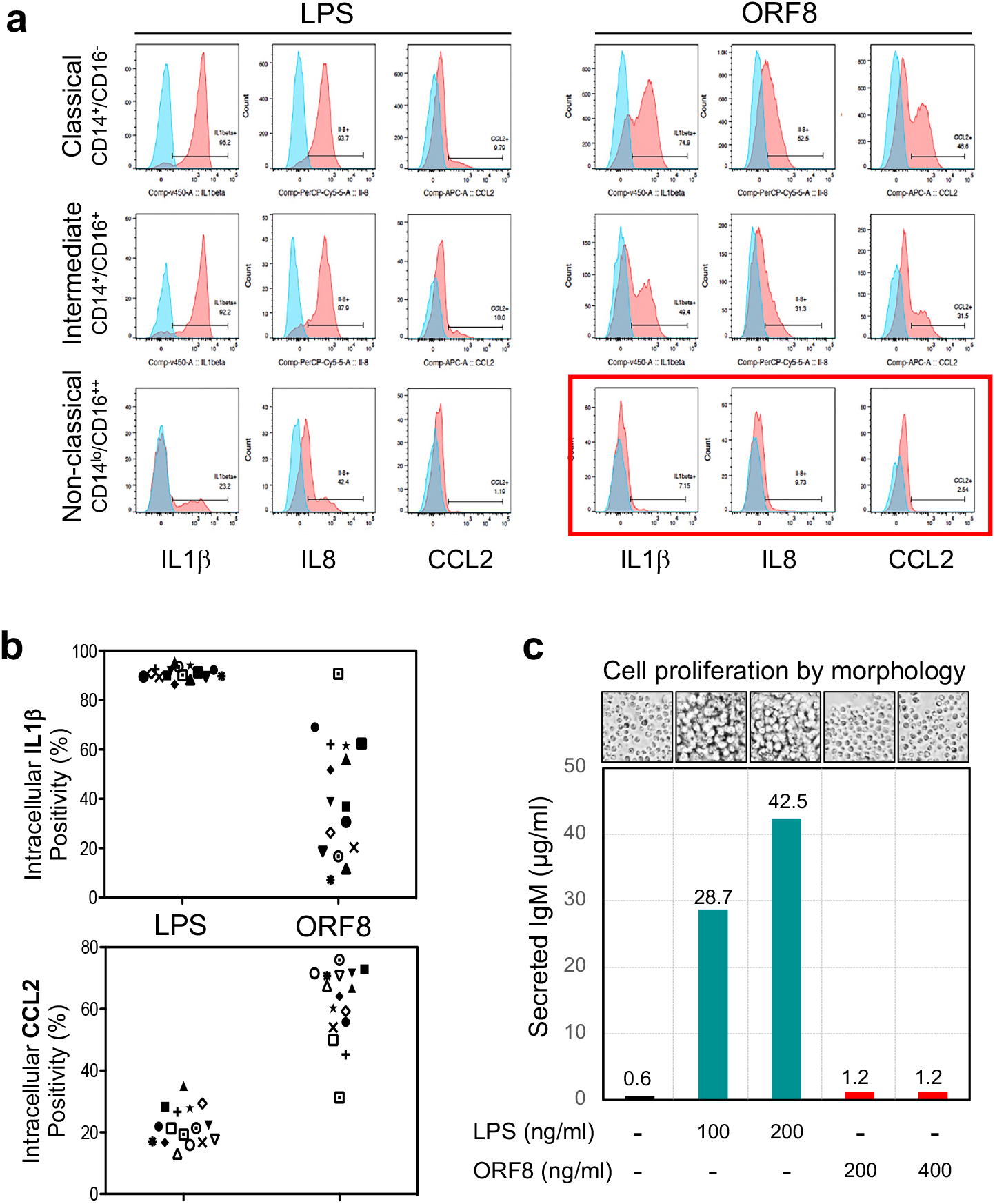
ORF8 is functionally different from LPS. **a.**Representative flow data showing LPS could activate multiple cytokine responses in all three monocyte subsets of PMBCs from all 15 healthy donors examined while ORF8 only activates CD14^+^ Monocytes but not CD14^low^/CD16^++^ non-classical monocytes (red boxed) suggesting LPS and ORF8 may target monocytes through different mechanisms. **b.** Intracellular cytokine flow cytometry data summary on 15 healthy donors upon treatment with ORF8 or LPS, suggesting that ORF8 and LPS are different in cytokine induction specificities. **c.** Mouse B cells responded differently to the treatment of ORF8 and LPS as measured by IgM production and the morphological features of cell proliferation status, demonstrating the lack of any detectable contaminating LPS activity in purified ORF8 protein.

### Secreted ORF8 directly binds to and activates the NLRP3-mediated inflammasome response

To decipher the mechanism by which ORF8 induces pro-inflammatory cytokines in CD14+ monocytes, using the Gene Set Enrichment Analysis (GSEA) and Kyoto Encyclopedia of Gene and Genome (KEGG) tools, we found that the SARS-CoV-2 infection pathway, NOD-like receptor signaling pathway, and NFkB pathway were among the top enriched, especially, mRNAs of lysosomal enzymes (LYZ, CTSD, CTSB, CTSS), inflammasomal protein NLPR3, and cytokines IL1β, IL8, CCL2 were among the top expressed (Fig. 3b).

We then examined the role of the NLRP3-mediated inflammasome pathway in ORF8-mediated cytokine production. Fig. 3d shows that both NLPR3 and IL1β proteins were induced in PBMCs upon ORF8 treatment, and such induction was diminished in NLPR3 knockdown cells (THP1-defNLRP3, InvivoGen) compared to the parental THP-1 cells, demonstrating the NLRP3 dependency of ORF8 mediated cytokine response in human monocytes (Fig. 3e).

To determine how ORF8 molecules enter CD14^+^ monocytes, we first incubated PBMCs with pure ORF8 protein on ice for one hour to saturate any potential ORF8 binding receptors on the monocyte surface. After extensive washes with ice-cold PBS to remove unbound ORF8 protein, the cells were then incubated at 37°C to initiate ORF8-mediated cell activation in the presence or absence of additional ORF8. Fig. 4a shows robust IL1β mRNA expression was observed only in cells exposed to additional ORF8 but not those only pre-incubated with ORF8, suggesting that ORF8 likely enters monocytes through a non-receptor-mediated process, such as phagocytosis. Complementarily, we asked if ORF8 would bind to TLR2, TLR4, CD14, or NLRP3 known to be involved in inflammasome activation. We incubated ORF8 protein-coated beads with monocyte lysates from two ORF8 responding and two ORF8 nonresponding healthy donors. Fig. 4b shows that neither TLR2, TLR4, nor CD14 were co-precipitated with ORF8 (CD14 data not shown). However, NLRP3 was readily detected in one of the ORF8-responders, suggesting that indeed ORF8 binds to NLRP3 in primary human monocytes.

To further dissect how NLRP3 binds to ORF8, we transiently transfected ORF8-expressing HEK293 cells with Flag-tagged NLRP3 constructs Flag-N3-FL (full-length NLRP3), or one of the three deletion mutants Flag-N3-N, Flag-N3-M, or Flag-N3-C (Fig. 4c).^22^ Cell lysates were then incubated with Strep-Tactin™ beads to pulldown Strep-tagged ORF8 and its binding proteins. In addition to NLRP3-FL (Fig. 4d), both Flag-N3-M, and Flag-N3-C but not Flag-N3-N also strongly co-precipitated with ORF8 suggesting both the middle NACHT domain and C-terminal LRR domain of NLRP3 each can independently bind to ORF8. These results clearly demonstrate inflammasome protein NLRP3 directly interacts with ORF8.

### ORF8 activates the NLRP3-mediated inflammasome pathways

The effect of LPS in inflammasomal activation has been well established; therefore, it was important to demonstrate that our purified ORF8 protein was free of LPS contamination. ORF8 and LPS were compared for their effects on IL1β, IL8, and CCL2 induction in three monocyte subsets (classical, intermediate, and non-classical) from 15 healthy donors. Fig 5a shows that all three subsets responded to LPS while only CD14^+^ (classical and intermediate) responded to ORF8 with variable induction of IL1β, IL8, and CCL2. These results show that ORF8 and LPS target monocytes with different cell type specificities. We then examined the responsiveness of CD14^+^ monocytes to LPS or ORF8 treatment using IL1β, IL8, and CCL2 expression as the readout, and we found that intracellular IL1β was detected in the majority (90.1%) of LPS-treated CD14^+^monocytes but only in a subpopulation (mean=38%) of ORF8 treated CD14^+^ monocytes. In contrast, CCL2 expression was detected only in a minor population (mean=22.1%) of LPS-treated CD14^+^ monocytes but in a larger population of ORF8-treated CD14+ monocytes (mean=63.7%) (Fig. 5b). These data clearly demonstrate that ORF8 and LPS also have different cytokine induction specificities. In addition, we also examined the cell proliferation and the IgM induction of mouse splenic B cells upon ORF8 and LPS treatment, and we found that the mouse splenic B cells did not proliferate, nor produced any polyclonal IgM 96 hours after ORF8 treatment, while LPS treated cells showed robust proliferation morphology and massive production of IgM (Fig. 5c). In the aggregate, our data clearly demonstrated that the cytokine inducing activity of ORF8 is not due to LPS contamination.

To examine the activation status of the canonical NFkB pathway necessary for the induction of inflammasomal proteins including NLPR3, Casp-1, and IL1β in ORF8 treated monocytes, we examined the GSEA analysis results and found that NFkB pathway components were transcriptionally enriched (NES=1.539662, FDR q-value=0) in ORF8 treated monocytes in cluster 5 (Fig. 6a, left panel). To validate these at the protein level, we treated THP-1 cells with ORF8 for 4 and 24 hours and determined their NFkB pathway protein expression. Fig. 6a (right panels) shows that the increased level of phospho-p65 and the conversion of p100 to p52 were readily detected 4 hours after ORF8 treatment. Similar results were also observed in LPS-treated cells. These data demonstrate that ORF8 stimulation activates the NFκB pathway triggering the activation of the inflammasome pathway in human monocytes.

**Figure 6.**
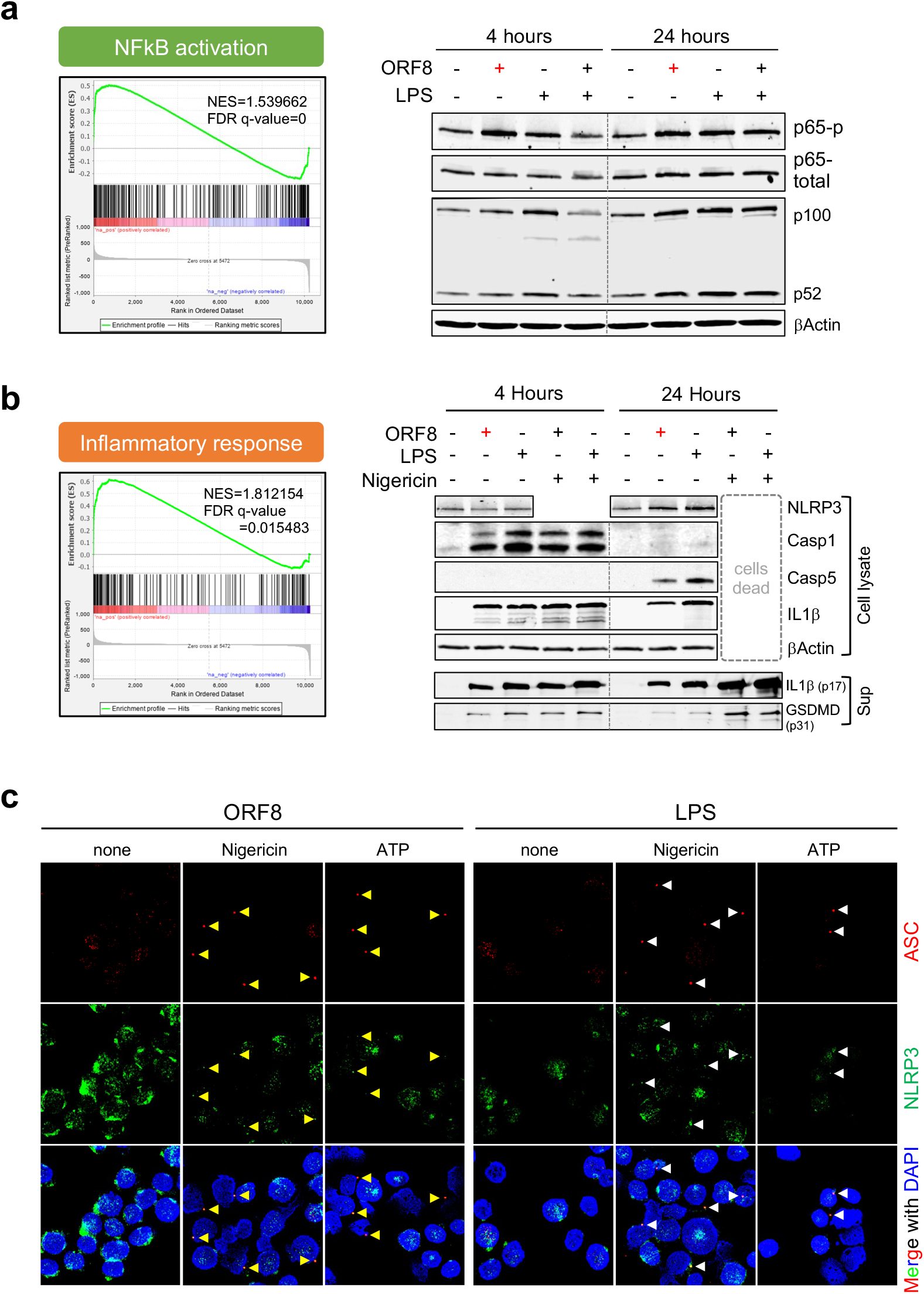
ORF8 induces proinflammatory cytokines through activation of NLRP3-mediated inflammasome pathways. **a.** Gene enrichment analysis of scRNA-Seq (left panel) and Western confirmation of key NFkB pathway molecules (right panels) upon stimulation of ORF8 for 4 or 24 hours. **b**. Gene enrichment analysis of scRNA-Seq (left panel) and Western confirmation of NLRP3, IL1β, Casp-1, and Casp-5 in THP-1 cell lysates and IL1β and GSDMD in the culture supernatants (right panels) upon stimulation of ORF8 for 4 or 24 hours. **c.** Immunofluorescence images show that, LPS and ORF8 activate ASC speck (arrowhead pointed) formation only when a second signal (nigericin or extracellular ATP) was present. Cells were first treated with 200 ng/ml ORF8 or 100 ng/ml LPS for 2.5 hours followed by 90 min treatment with 5 μM Nigericin or 5 μM ATP before cell harvest and staining for immunofluorescence.

Similarly, the GSEA analysis data revealed the mRNAs of inflammasome pathway components were also significantly enriched (NES=1.812154, FDR q-value =0.015483) (Fig. 6b, left panel). We then treated THP-1 cells with ORF8 or LPS alone or in combination with nigericin, a potent agent that provides a second signal for the canonical inflammasomal pathway activation, for 4 or 24 hours. The cell lysates and culture supernatants were analyzed by Western blotting for the expression of NLPR3, Caspase-1, Caspase-5, IL1β, and the pore-forming fragment of gasdermin D (GSDMD) proteins. We found that ORF8 induces the inflammasome pathway by 1) activating the production of IL1β without requiring a separate priming step; 2) signaling through the non-canonical pathway using Casp-1 (early hours) followed by switching to Casp-4/5; 3) maintaining cell viability without triggering pyroptosis unless the second signal such as nigericin is present which rapidly initiates pyroptosis (Fig. 6b, right panel). Our data suggest that ORF8 and LPS may represent a family of pyrogens by triggering a non-canonical inflammasomal pathway leading to the production and release of IL1β without activating GSDMD to the level necessary for pyroptosis, a process called hyperactivation for LPS.^23,24^ However, ORF8 can also serve as a priming agent for the canonical inflammasome pathway to evoke a prompt pyroptosis in the presence of second signal molecules such as extracellular ATP from damaged cells.

To validate the dual roles of ORF8 in the activation of canonical and non-canonical inflammasome pathways, we examined ASC speck formation in ORF8- or LPS-treated THP-1 cells by immunofluorescence. As shown in Fig. 6c, ORF8 alone was insufficient to induce ASC speck formation. However, distinct pre-nuclear ASC specks were readily detected when nigericin or extracellular ATP was also present. In addition, the NLPR3 protein was also co-localized with ASC protein to the specks (Fig. 6c), ASC speck formation always precedes cell pyroptosis in our system, and ORF8 is slightly more robust than LPS in mediating ASC formation in the presence of nigericin or extracellular ATP. The results confirm the dual roles of ORF8 in the activation of the non-canonical inflammasome pathway when the second signal is absent, and the canonical inflammasome pathway by priming the cells when the second signal is present. These data further suggest that the mode of action of ORF8 is dependent on the presence or absence of a second signal such as extracellular ATP released from damaged cells/tissues nearby.

### Blood ORF8 protein levels correlate with the mortality and disease course of severe patients with COVID-19

Having demonstrated that ORF8 is secreted as a glycoprotein and capable of inducing inflammatory cytokine responses *in vitro*, we then asked if ORF8 is also secreted and glycosylated in blood from patients infected with SARS-CoV-2. By analyzing serum samples collected from patients within 4-7 days of COVID-19 diagnosis using Western blotting, we found that ORF8 protein was readily detected at various levels in 92% (23/25) of newly infected patients. Interestingly, two other major secreted proteins NSP9 and NSP10 were not detected in the same samples suggesting not all the secreted SARS-CoV-2 proteins are present at detectable levels in blood circulation. To determine the glycosylation status of the serum ORF8, we compared the sizes of ORF8 from patients and HEK293 lysate. Our results (Fig. 7A) show that patient serum ORF8 is indeed glycosylated when compared to the ORF8 from HEK293 lysate which has an extra Strep-II tag (28 aa) and possibly an uncleaved signal sequence (16 aa). Due to the lack of a quantitative assay, we were unable to determine the absolute quantity of ORF8 in patient samples; however, our semi-quantitative Western blot could readily detect ORF8 in as little as four microliters of patient serum suggesting the protein is present at significant levels in patient blood at the onset of COVID-19.

**Figure 7.**
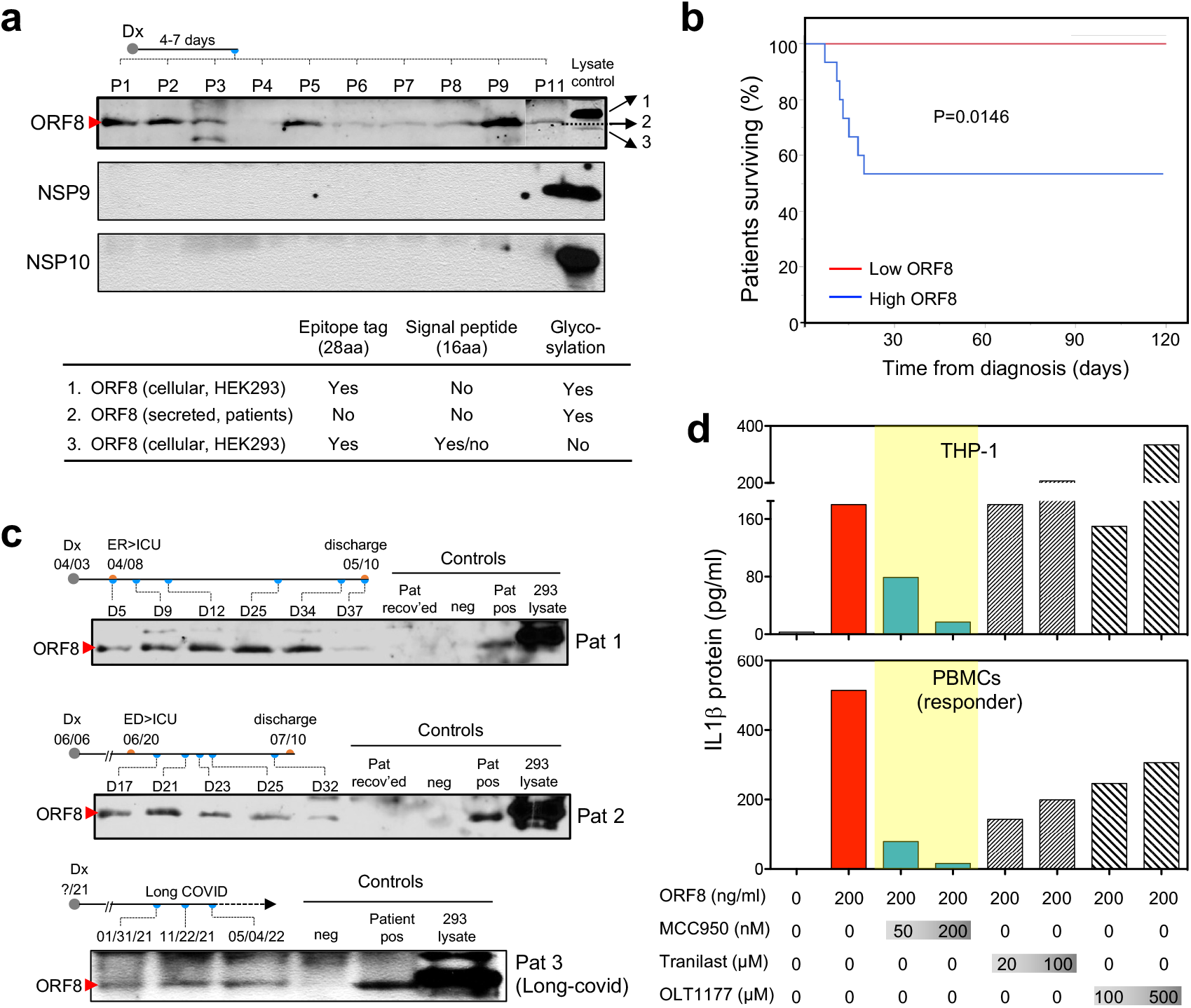
The levels of ORF8 protein in sera of COVID-19 patients correlate with the outcome and disease course, and the ORF8 pathway is targetable with NLRP3 inhibitors. **a.** Representative Western blot images showing ORF8 protein but not NSP9 or NSP10 were detectable in as little as 4.0 μl of serum samples from newly infected COVID-19 patients. ORF8 protein in patient sera shown as glycosylated by size estimation; **b.** Kaplan-Meier curve showing fatality in hospital patients is associated with higher serum level of ORF8; **c.** Western images showing ORF8 levels at different time points correlate with disease course in two patients with prolonged clinical courses, and in one patient with long-covid. **d.** Targetability of ORF8 mediated IL1β expression with various NLRP3 inhibitors in THP1 cells and human primary PBMCs. The data shown are representative of two or three independent experiments. Abbreviations: Dx = diagnosis; ED = emergency department; ICU = intensive care unit

Given its activity in inducing inflammatory cytokines *in vitro*, and its significant presence in the blood of newly infected patients, we then asked if the levels of ORF8 protein would correlate with disease severity and outcome. To that end, we examined the correlation between blood ORF8 levels and patients’ survival outcomes in our cohort of 25 hospitalized patients. Fig. 7b shows that after 120 days of follow-up, all seven fatalities were exclusively associated with the ORF8-high group (scored 2+ or 3+ on Western blot) while all patients in the ORF8-low group (scored 0 or 1+) had mild symptoms and quick recovery without any events of death. These results demonstrate a clear correlation between the ORF8 load and disease severity in newly infected patients. We then monitored the ORF8 levels at various time points during the disease course in two patients with prolonged COVID-19, and one patient with long covid lasting more than 17 months. As demonstrated in Fig. 7c, two patients with high blood ORF8 levels had persistent severe symptoms requiring treatment in the intensive care unit (ICU), and their blood ORF8 levels diminished by the time they were recovered and discharged from the hospitals. The long covid patient with lingering symptoms throughout remained ORF8 positive in all three serial blood samples collected over 16-month period (Fig. 7c). These results suggest that ORF8 level is prognostic for COVID-19 outcome, further supporting our hypothesis that the ORF8 protein is a pathogenic cause of severe COVID-19 in patients.

### ORF8-mediated cytokine induction is targetable by select NLRP3 inhibitors

Given its causative role in COVID-19 pathogenesis, we then tested the targetability of ORF8 mediated cytokine response using three investigational NLRP3 inhibitors - MCC950, Tranilast, and OLT117 (Dapansutrile). As shown in Fig. 7d, the NLRP3 inhibitor MCC950 showed effective inhibition on the production of IL1β in both THP-1 cells and PBMCs (blue bars), while Tranilast and OLT117 (hatched bars) exhibited moderate to marginal effect in PBMCs but no effect in THP-1 cells. Our results suggest that ORF8 mediated IL1β production can be effectively inhibited by the NLRP3 inhibitor MCC950 at a nanomolar dose range, and possibly by other classes of inhibitors as well, demonstrating that targeting the ORF8/NLRP3 axis is a promising strategy for treating patients with symptomatic COVID-19.

## DISCUSSION

Despite the success in the development and implementation of effective vaccines to prevent SARS-CoV-2 infection, COVID-19 remains a major challenge for many reasons. These include 1) a significant population remains unvaccinated, 2) new variants keep emerging and evading current vaccine protection, 3) patients with compromised immunity including those with blood cancers on immunosuppressive medications are poorly protected, ^25,26^ and more importantly, 4) many previously infected patients have developed lingering symptoms or long Covid. Therefore, better therapies and clinical management tools for COVID-19 are urgently needed.

SARS-CoV-2 accessory protein ORF8, a 121 amino acid protein, is the least conserved protein in the beta-coronavirus family.^27,28^ Here we report that ORF8 is a major secreted viral glycoprotein *in vitro* and in COVID-19 patients. Our data, summarized in Supplemental Fig. 7, support the concept that the ORF8 protein secreted from locally infected cells in the lung and released to the bloodstream then stimulates circulating CD14^+^ monocytes to initiate systematic cytokine responses. This is consistent with findings by others that monocytes are the single most affected WBC subset in the blood by SARS-CoV-2 infection.^29^ It has been reported that about 6% of monocytes were infected with the virus in COVID-19 patients through CD16-mediated uptake of antibody-opsonized SARS-CoV-2 virus. Whether these cells were truly infected or simply engulfed with infected cells/debris remains to be seen since no live virus was detected.^30^ Nevertheless, these monocytes may represent a parallel and complementary process to the ORF8-mediated inflammatory pathway described herein. Our intracellular IL1β data showed a mean of 43% (range, 7.2% - 90.7%) of monocytes that are ORF8 responsive, suggesting that the ORF8-mediated process dominates. We conclude that secreted ORF8 is a key disease-causing viral factor and can be targeted by NLRP3 inhibitors, offering a potential new treatment option.

NLRP3-mediated inflammasome activation is a critical part of the innate immune response. However, the precise mechanisms for different pathogens have not been fully delineated. Our study indicated that ORF8 may enter monocytes through a non-receptor mediated ORF8 internalization, followed by lysosomal action involving lysozyme (LYZ) and cathepsin proteases (CTSB, CTSD). ORF8 then may bind to the NACHT and/or LRR domain to activate NLRP3 and recruit Casp-1 (early) and Casp-4/5 (later). It is not known what the roles of these two different bindings play, and if they are important in pathway choosing (canonical vs. non-canonical). It is also interesting that, like LPS, ORF8 induces the expression/activation of Caspase-1 early (hour 4) but switches to Casp-4/5 later (at hour 24) even when nigericin is present (data not shown). Our data shows that the mode of action of ORF8 is readily changed by the presence of the second signal leading to the switch from non-canonical to canonical inflammasomal response. Therefore, it is conceivable that, in patients with COVID-19, the inflammation may be aggravated by the additional tissue/cell damages that release ATP, leading to the activation of an ORF8-mediated canonical inflammasome pathway and causing pyroptosis.

Due to the lack of effective COVID-19 animal models, it is challenging to genetically test the pathogenic role and targetability of the ORF8 protein *in vivo*. However, a SARS-CoV-2 variant (Δ382) found in Taiwan and Singapore with a complete loss of the ORF8 gene serves as a tailor-made natural genetic model that validates our findings.^31,32^ Young et al. analyzed a cohort of 92 patients infected with the wildtype virus and 29 patients infected with the Δ382 variant and found that patients infected with the wildtype SARS-CoV-2 exhibited much higher levels of pro-inflammatory cytokines than those infected with the Δ382 variant. Furthermore, no patients with the Δ382 variant required supplemental oxygen whereas it was required for 26 (28%) patients infected with wild-type virus. These observations provide yet another line of evidence supporting the link between ORF8 and COVID-19 disease severity, and an *in vivo* genetic proof-of-principle for targeting ORF8 to prevent severe COVID-19 infection. Additionally, various clinical studies and trials have shown that the IL1 receptor antagonist anakinra significantly reduced the mortality risk in hospitalized patients with moderate to severe inflammation symptoms.^33 34,35^ The data from our present study suggests that targeting NLRP3 or ORF8 upstream of the IL1 receptor would provide greater therapeutic benefit to patients with severe COVID-19 symptoms, and could also provide prophylactic benefit to patients at risk of developing severe COVID-19 infection.

## ACKNOWLEDGEMENT

We thank The Mayo Clinic Genome Analysis Core, Bioinformatics Core, Microscopy and Cell Analysis Core, and the CMM Electron Microscopy Facility of the University of California San Diego (supported by NIH equipment grant 1S10OD023527-01) for their expert service, Mayo Clinical Blood Bank for providing fresh leukocyte cones of healthy donor blood, and the Mayo Clinic Center for Individualized Medicine for providing access to their biobank data. We also thank the University of Iowa/Mayo Clinic Lymphoma SPORE (NIH/NCI grant P50 CA97274) and the Predolin Foundation Biobank for sharing some of their resources for this project.

## AUTHOR CONTRIBUTION

XW and TEW designed the experiments, analyzed, and interpreted results, and wrote the manuscript; XW, MKM, GJR, KEN, KAG, XT, TLW, VT, AW, and MJS conducted experiments and analyzed data; XW, JPA, YY, XT, SD, and MJS analyzed data and generated figures; ADB, JPA, GR, JP, SMA, VB, HD, and TEW provided experiment samples; JPA, GJR, JP, TLW, SMA, MJS discussed and interpreted results; all authors read and approved the manuscript.

