## Supplemental (methods-Tables-Figures) for "Secreted ORF8 is a pathogenic cause of severe COVID-19 and is potentially targetable with select NLRP3 inhibitors"

**Included:**

Additional Materials and Methods

Supplemental Tables (4)

Supplemental Figures (7)

**Materials and Methods (supplemental):**

*Single-cell RNA sequencing of PBMCs treated with ORF8 or control medium*

Three million PMBCs were treated with either media control- or ORF8-CM for three days; cells were collected and enriched for viable cells to greater than 90% using the dead cell removing kit (Miltenyi Biotec, Cat#130-090-101), submitted for cell capturing, and single-cell RNA sequencing. The cell suspension and Chromium Single Cell 3′ v2 library master mix (10x Genomics), gel beads, and partitioning oil were added to a Chromium Single Cell A chip which was then loaded into the Chromium Controller for capturing 4000 single cells and subsequent cDNA synthesis. The resulting cDNA libraries were pooled and sequenced as 100x2 paired-end reads on an Illumina HiSeq 4000 using HiSeq 3000/4000 sequencing kit.

*Single-cell RNA sequencing Data Analysis*

We used the Seurat package (v4.0.1) to perform integrated analyses of single cells. Genes expressed in less than 3 cells, cells that expressed less than 200 genes, and more than 40% which were mitochondria genes were excluded for downstream analysis in each sample. For scRNA-Seq analysis, we followed the Seurat integration workflow and the comparative analysis workflow. Each dataset was normalized and the top 2000 Highly Variable Genes (HVGs) across cells were selected. The datasets were integrated based on "anchors" identified between datasets before Principal Component Analysis was performed to do a linear dimensional reduction. Shared Nearest Neighbor (SNN) Graph was constructed to identify clusters in the low-dimensional space (top 50 statistically significant principal components). Enriched marker genes in each cluster conserved across all samples were identified, and differentially expressed genes between two selected conditions were detected using the default Wilcoxon Rank Sum test at the cluster level. Both GSEA and KEGG pathway enrichment analyses were performed.

*Detection of SARS-CoV-2 protein secretion*

To detect protein secretion, HEK293 cells expressing the protein of interest were seeded at 70% confluency in a T25 flask in 4ml of DMEM/10% FCS. After 48 hours of culture, the supernatants were collected and cleared for cells and debris by centrifugation at 3000 rpm for 5 min. Four mL of cleared supernatants were then incubated overnight with 40ul of a 50% slurry of StrepTactin (or anti-HA for SPK protein) beads. The beads were then washed twice with PBS and then resuspended in 40 ml Laemmli sample buffer with 5% BME. The amount of sample equivalent to 1ml of culture supernatant was loaded into a 4-20% gradient gel followed by western blotting per routine probing with anti-Strep II antibody at 1:1000 dilution (#71590, Millipore) and developed on a Li-Cor scanner.

*Expression and purification of ORF8 from HEK293F cell supernatant*

Transfected HEK293F cells were grown for 3-4 days at a cell density below $6x10^6$ cells per ml. Cells were pelleted by centrifugation and supernatant containing secreted ORF8 was supplemented with a Complete-EDTA free protease inhibitor cocktail (Roche), 1mM BME, and 1 mL of biotin blocking solution (BioLock). The media were loaded on a 5 mL StrepTactin Sepharose High-Performance column (GE Healthcare) in PBS buffer and eluted with a linear gradient of 0-50% 5 mM Biotin in PBS. Fractions containing ORF8 were pooled, concentrated by ultrafiltration (Amicon), and then purified using a Superdex 200 Increase 10/300 GL (Cytiva) in PBS, followed by anion exchange chromatography using a Source 15Q 4.6/10 column (Cytiva) with a gradient of 0–500 mM NaCl in 20mM Tris pH 7.5. Fractions containing ORF8 were pooled and concentrated by ultrafiltration and stored at 4°C.

*Expression and purification of ORF8 from E. Coli*

DNA encoding SARS CoV-2 ORF8 codon-optimized for expression in *E. coli* was cloned into pMCSG7 [1] and transformed into Rosetta2 cells (EMD), and protein expression was induced with 100 μM IPTG overnight at 15°C. *E. coli* culture was pelleted, lysed, and protein purified as described previously [2], except that cells were lysed by sonication with a Branson sonicator in place of Dounce homogenization. The concentrated, refolded protein was loaded on a Superdex S75 size-exclusion column equilibrated in PBS. Fractions containing ORF8 were pooled and concentrated by ultrafiltration (Amicon). The His-tag was removed by incubation with TEV protease. Untagged ORF8 was run on a Source 15Q 4.6/10 anion exchange column (Cytiva) equilibrated in wash buffer (20 mM Tris pH 7.5) and eluted with a linear gradient of 0-100% 500 mM NaCl in 20 mM Tris pH 7.5. Monomer and dimer peaks were pooled and concentrated.

*Detection of ORF8 protein in culture supernatants with Luminex method*

To detect cytokines in the culture supernatant we used two methods – the mRNA results from qPCR and a custom Luminex detection kit for IL1β, IL6, IL8, and TNFa) (R&D systems) or a custom Luminex cytokine detection kit for IL1β, IL6, IL8, IL18, CCL2, and TNFα (ThermoFisher). The mRNA results from qPCR of the same experiment were compared with the protein data from Luminex.

*Immunofluorescence detection of ASC specks*

Thp-1 cells were treated with ORF8 at 200 ng/ml or LPS at 100 ng/ml for 2.5 hours followed by adding 0 or 5 ug/ml Nigericin for an additional 90 min (total of 4 hours), the cells amounted to slides by cytospin, fixed with 4% PFA for 10 min, and stained and imaged for ASC and NLRP3 according to previously described staining and imaging methods.[3]

*Intracellular cytokine measurement.*

Two million PBMCs were activated with ORF8 or LPS for 18.5 hours, and Golgi-Plug (contains Brefeldin A) (Cat # 51-2301KZ, BD Pharmingen) was added at 1:1000 for additional 5.5 hours of incubation. The cells were then collected and stained with cell lineage surface markers (CD3, CD19, CD14, CD16, CD11c, CD56, HLA-Dr) and FVD for live/dead gating followed by staining intracellular IL1β, IL8, IL6, and CCL2 upon fix and permeabilization using a BD Cytofix/Cytoperm fixation/Permeabilization Kit (Cat # 554714). All antibodies used are listed in Supplemental Table 4. The flow cytometry 13-color detections were run on an LSRFortessa X-20 (BD). The Data were analyzed using Flowjo software V10.

**REFERENCES (of the section)**


1. Stols L, Gu M, Dieckman L, Raffen R, Collart FR, Donnelly MI. A new vector for high-throughput, ligation-independent cloning encoding a tobacco etch virus protease cleavage site. *Protein Expr Purif*. 2002;25(1):8-15.

2. Flower TG, Buffalo CZ, Hooy RM, Allaire M, Ren X, Hurley JH. Structure of SARS-CoV-2 ORF8, a rapidly evolving immune evasion protein. *Proc Natl Acad Sci U S A*. 2021;118(2).

3. Wu X, Platt JL, Cascalho M. Dimerization of MLH1 and PMS2 limits nuclear localization of MutLalpha. *Mol Cell Biol*. 2003;23(9):3320-3328.

### Supplemental Tables

**Supplemental Table 1.** SARS-CoV-2 constructs used in these study

| Addgene ID | Construct Name |
|---|---|
| 141367 | pLVX-EF1alpha-SARS-CoV-2-**NSP1**-2xStrep-IRES-Puro |
| 141368 | pLVX-EF1alpha-SARS-CoV-2-**NSP2**-2xStrep-IRES-Puro |
| 141369 | pLVX-EF1alpha-SARS-CoV-2-**NSP4**-2xStrep-IRES-Puro |
| 141371 | pLVX-EF1alpha-SARS-CoV-2-**NSP5**-C145A-2xStrep-IRES-Puro |
| 141373 | pLVX-EF1alpha-SARS-CoV-2-**NSP7**-2xStrep-IRES-Puro |
| 141374 | pLVX-EF1alpha-SARS-CoV-2-**NSP8**-2xStrep-IRES-Puro |
| 141375 | pLVX-EF1alpha-SARS-CoV-2-**NSP9**-2xStrep-IRES-Puro |
| 141376 | pLVX-EF1alpha-SARS-CoV-2-**NSP10**-2xStrep-IRES-Puro |
| 141378 | pLVX-EF1alpha-SARS-CoV-2-**NSP12**-2xStrep-IRES-Puro |
| 141379 | pLVX-EF1alpha-SARS-CoV-2-**NSP13**-2xStrep-IRES-Puro |
| 141380 | pLXV-EF1alpha-2xStrep-SARS-CoV-2-**NSP14**-IRES-Puro |
| 141381 | pLVX-EF1alpha-SARS-CoV-2-**NSP15**-2xStrep-IRES-Puro |
| 141347 | pBOB-CAG-SARS-CoV2-**SPK**-HA |
| 141383 | pLVX-EF1alpha-SARS-CoV-2-**ORF3a**-2xStrep-IRES-Puro |
| 141384 | pLXV-EF1alpha-2xStrep-SARS-CoV-2-**ORF3b**-IRES-Puro |
| 141385 | pLVX-EF1alpha-SARS-CoV-2-**ENV**-2xStrep-IRES-Puro |
| 141386 | pLVX-EF1alpha-SARS-CoV-2-**MEM**-2xStrep-IRES-Puro |
| 141387 | pLVX-EF1alpha-SARS-CoV-2-**ORF6**-2xStrep-IRES-Puro |
| 141388 | pLVX-EF1alpha-SARS-CoV-2-**ORF7a**-2xStrep-IRES-Puro |
| 141389 | pLXV-EF1alpha-2xStrep-SARS-CoV-2-**ORF7b**-IRES-Puro |
| 141390 | pLVX-EF1alpha-SARS-CoV-2-**ORF8**-2xStrep-IRES-Puro |
| 141391 | pLVX-EF1alpha-SARS-CoV-2-**NUC**-2xStrep-IRES-Puro |
| 141394 | pLVX-EF1alpha-SARS-CoV-2-**ORF10**-2xStrep-IRES-Puro |
| 75127 | pcDNA3-N-Flag-**NLRP3-FL** |
| 75137 | pcDNA3-N-Flag-**NLRP3 1-90** |
| 75140 | pcDNA3-N-Flag-**NLRP3 91-710** |
| 75141 | pcDNA3-N-Flag-**NLRP3 711-1034** |

**Supplemental Table 2.** Top 20 enriched genes expressed in cells of cluster 5

| Top genes | Function | ORF8 Avg_log2FC |
|---|---|---|
| IL8 | Cytokine/chemokine | 5.14 |
| CCL2 | Cytokine/chemokine | 4.92 |
| IL1B | Cytokine/chemokine | 3.53 |
| | **Monocyte-related** | |
| CD14 | Monocyte cell marker | 3.66 |
| CD68 | Monocyte cell marker | 3.38 |
| FCER1G | IgE Fc receptor | 3.77 |
| | **Lysosome and inflammation-related** | |
| LYZ | Lysosomal protease | 5.07 |
| MMP9 | Tissue remodeling & cell mobilization | 5.01 |
| S100A9 | Inflammatory response & Cytokine secretion | 4.55 |
| S100A8 | Inflammatory response & Cytokine secretion | 3.82 |
| FTH1 | Ferritin heavy chain | 4.39 |
| FTL | Ferritin light chain | 3.77 |
| CTSD | Lysosomal A1 family peptidase | 3.82 |
| CTSB | Lysosomal C1 family peptidase | 3.69 |
| CTSS | Lysosomal C1 family peptidase | 3.59 |
| ACP5 | Elevated during inflammation | 3.68 |
| AIF1 | Elevated during inflammation | 3.53 |
| FUCA1 | Lysosomal fucose-glycoprotein degradation | 3.58 |
| IFI30 | Lysosomal thiol reductase | 4.50 |
| CYBB | Microbicidal oxidase of phagocytes | 3.79 |

**Supplemental Table 3.** qPCR Primers used for detection of cytokine expression

| Cytokine | Primer name | Primer sequence |
|---|---|---|
| IL1b | IL1b-qF | CCACAGACCTTCCAGGAGAATG |
| | IL1b-qR | GTGCAGTTCAGTGATCGTACAGG |
| IL6 | IL6-qF | AGACAGCCACTCACCTCTTCAG |
| | IL6-qR | TTCTGCCAGTGCCTCTTTGCTG |
| IL8 | IL8-qF | GAGAGTGATTGAGAGTGGACCAC |
| | IL8-qR | CACAACCCTCTGCACCCAGTTT |
| CCL2 | CCL2-qF | AGAATCACCAGCAGCAAGTGTCC |
| | CCL2-qR | TCCTGAACCCACTTCTGCTTGG |
| IL10 | IL10-qF | TCTCCGAGATGCCTTCAGCAGA |
| | IL10-qR | TCAGACAAGGCTTGGCAACCCA |
| CCL3 | CCL3-qF | ACTTTGAGACGAGCAGCCAGTG |
| | CCL3-qR | TTTCTGGACCCACTCCTCACTG |
| CCL5 | CCL5-qF | CCTGCTGCTTTGCCTACATTGC |
| | CCL5-qR | ACACACTTGGCGGTTCTTTCGG |
| IP10 | IP10-qF | GGTGAGAAGAGATGTCTGAATCC |
| | IP10-qR | GTCCATCCTTGGAAGCACTGCA |
| TNFa | TNFa-qF | CTCTTCTGCCTGCTGCACTTTG |
| | TNFa-qR | ATGGGCTACAGGCTTGTCACTC |
| GM-CSF | GMCSF-qF | GGAGCATGTGAATGCCATCCAG |
| | GMCSF-qR | CTGGAGGTCAAACATTTCTGAGAT |

##### Supplemental Table S4. Key antibodies used in this study

| Antibody | Source or Fluorophore | Clone | Vendor | Cat# |
|---|---|---|---|---|
| **For Western Blotting** | | | | |
| ASC (B-3) | mouse | B-3 | Santa cruz | sc-514414 |
| Caspase-5 | mouse | D3G4W | Cell Signaling | 46680 |
| Cleaved Caspase-1 (Asp297) | rabbit | D57A2 | Cell Signaling | 4199S |
| FLAG | mouse | M2 | Sigma | F3165 |
| Gasdermin D | rabbit | E8G3F | Cell Signaling | 97558 |
| IL-1 beta | rabbit | EPR21086 | Abcam | ab216995 |
| NF-kB p65 | mouse | L8F6 | Cell Signaling | 6956 |
| NF-kB2 p105/p50 | rabbit | D7H5M | Cell Signaling | 12540 |
| NLRP3 | rabbit | D4D8T | Cell Signaling | 15101S |
| NSP10 SARS-CoV-2 | mouse | 1049919 | R&D | MAB11049-SP |
| NSP9 SARS-CoV-2 | rabbit | EPR24855-25 | Abcam | ab284037 |
| ORF8 SARS-CoV-2 | rabbit | | MyBioSource | MBS3014575 |
| pNFKB p65 (Ser536) | rabbit | 93H1 | Cell Signaling | 3033 |
| Strep-Tag® II | mouse | GT661 | Millipore/Novagen | 71590-3 |
| β-Actin | mouse | C4 | Santa Cruz | sc4778 |
| **For Flow Cytometry** | | | | |
| CD19 | BV650 | SJ25-C1 | BioLegend | 363025 |
| CD3 | FITC | OKT3 | BioLegend | 317306 |
| CD14 | APC-Cy7 | M5E2 | BioLegend | 301820 |
| CD16 | BUV 496 | 3G8 | BD | 564653 |
| CD11c | PE-cy7 | B-ly6 | BD | 561356 |
| CD56 | PE-dazzle | HCD56 | BioLegend | 318347 |
| CD5 BV605 | BV605 | L17F12 | BD | 742550 |
| HLA-DR | BV785 | GF6-6 | BD | 564041 |
| Ghost dye viability dye | VIOLET 510 | | Tonbo | 13-0870-T100 |
| IL-1β | BV421 | BV421 | BD Pharmingen | 567791 |
| IL-6 | PE | MQ1-6A3 | BD Pharmingen | 559331 |
| Il-8 | PerCP-eFlour710 | 8CH | Invitrogen | 46-8088-42 |
| CCL2 (MCP-1) | APC | 5D3-F7 | Invitrogen | 17-7099-81 |

### Supplemental Figures

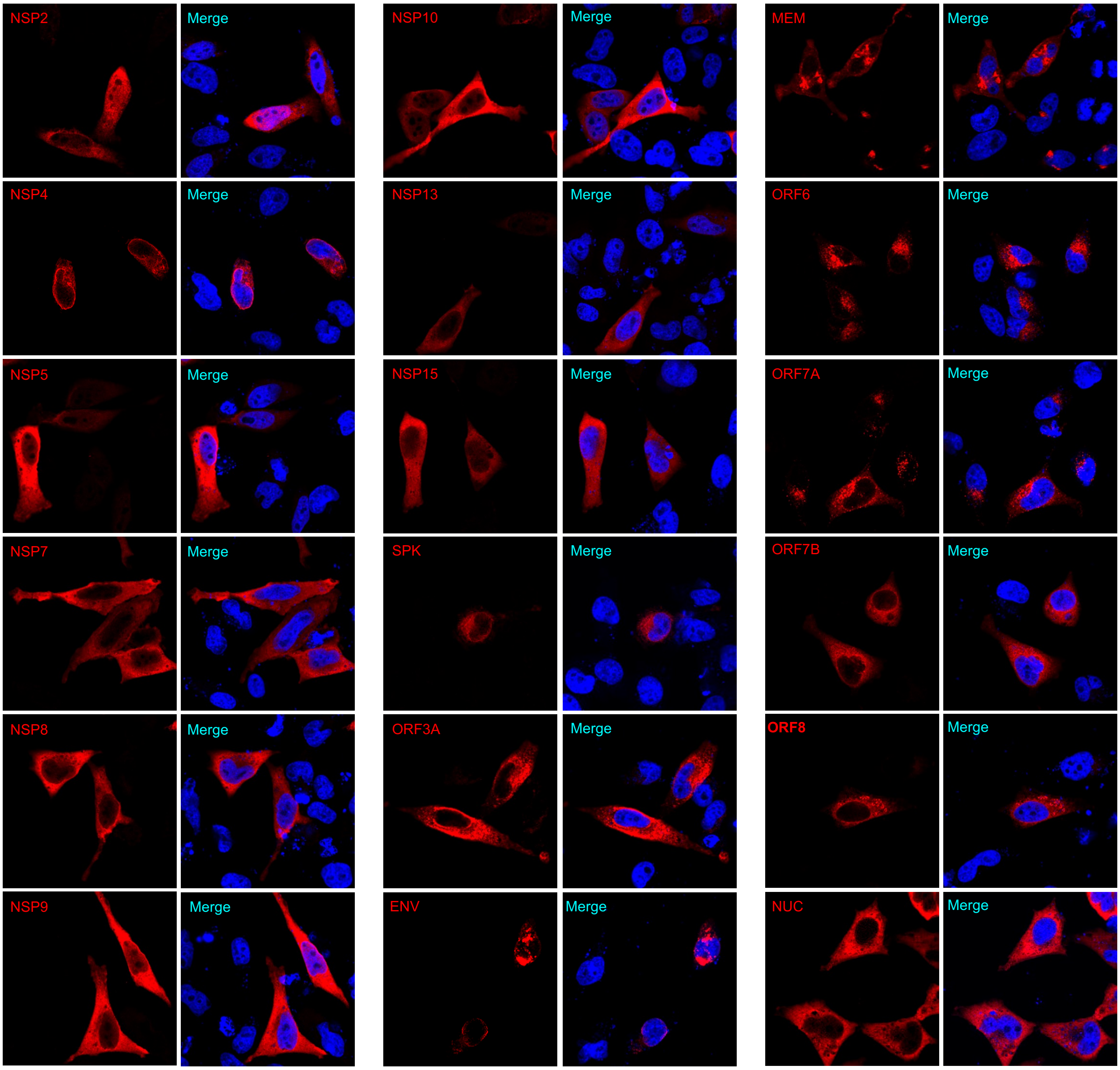


**Supplemental Figure 1. Intracellular localization of major proteins encoded by SARS-CoV-2 expressed in HeLa cells.** Confocal images of HeLa cells transiently transfected with respective SARS-CoV-2 constructs for 24 hours, stained with anti-Strep II antibody or anti-HA for Spike proteins (Red) and counterstained with DAPI (blue).

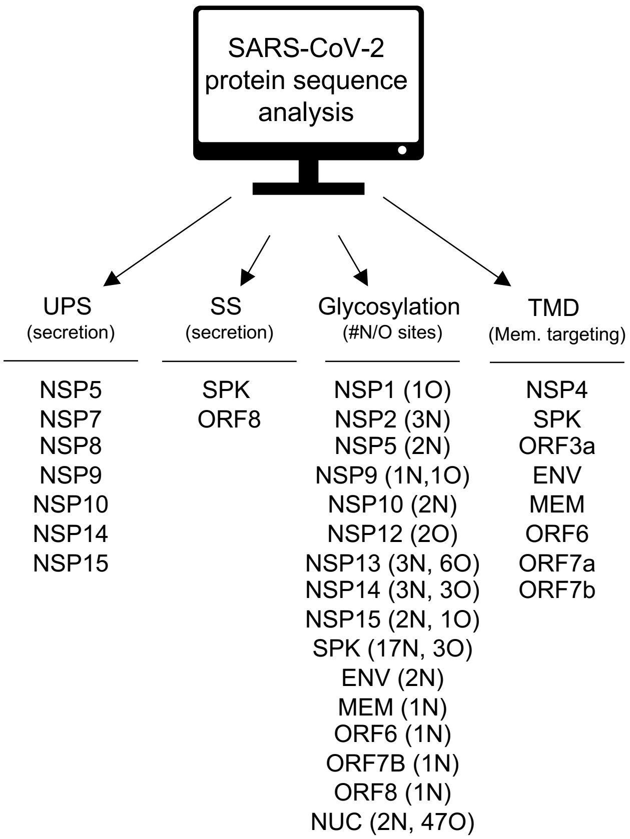


**Supplemental Figure 2.** ***In silico*** **protein sequence analysis of SARS-CoV-2 gene products.** Online tools used were Outcyte 1.0 server (http://www.outcyte.com/analyse/) for UPS analysis, SignalP 5.0 server (http://www.cbs.dtu.dk/services/SignalP/) for SS analysis, NetNGlyc 1.0 Server (http://www.cbs.dtu.dk/services/NetNGlyc/) for N-glycosylation site prediction, NetOGlyc 4.0 Server (http://www.cbs.dtu.dk/services/NetOGlyc/) for GalNAc O-glycosylation site prediction, and TMHMM 2.0 (http://www.cbs.dtu.dk/services/TMHMM-2.0/) for TMD analysis.

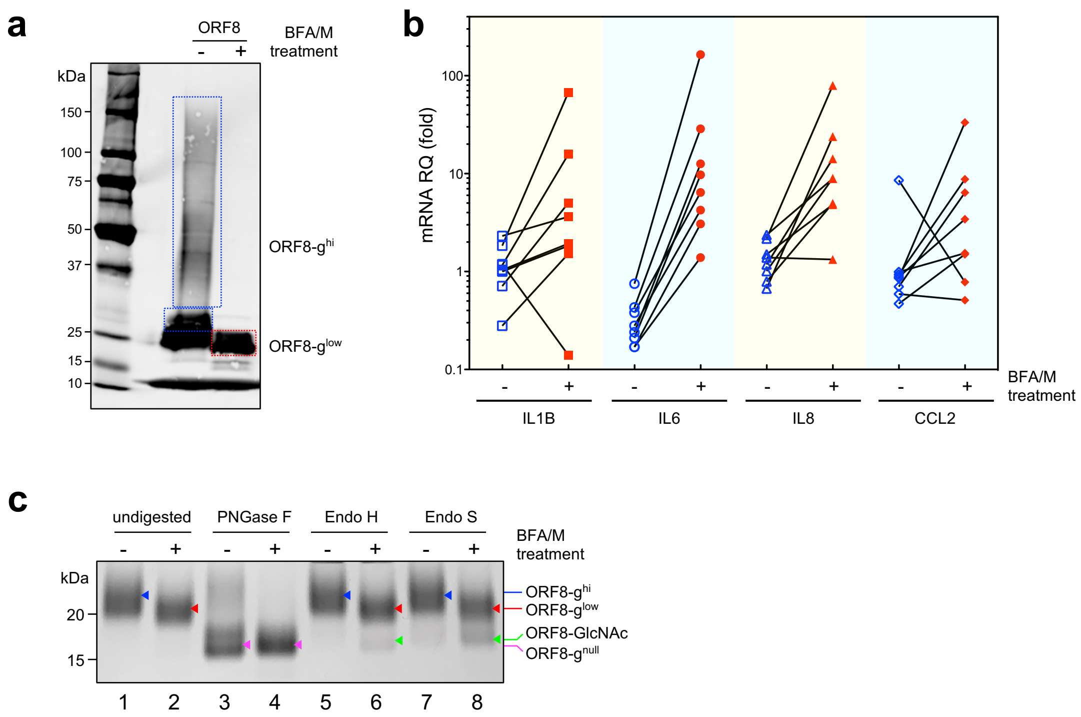


**Supplemental Figure 3. Inhibition of Golgi function modifies the glycosylation and cytokine induction activity of secreted ORF8 protein. a.** ORF8 protein mobility is affected by the treatment of Golgi inhibitor BFA/M; **b.** ORF8 secreted from BFA/M treated cells is more active in cytokine induction; **c.** Characterization of purified ORF8 using various glycosylases.

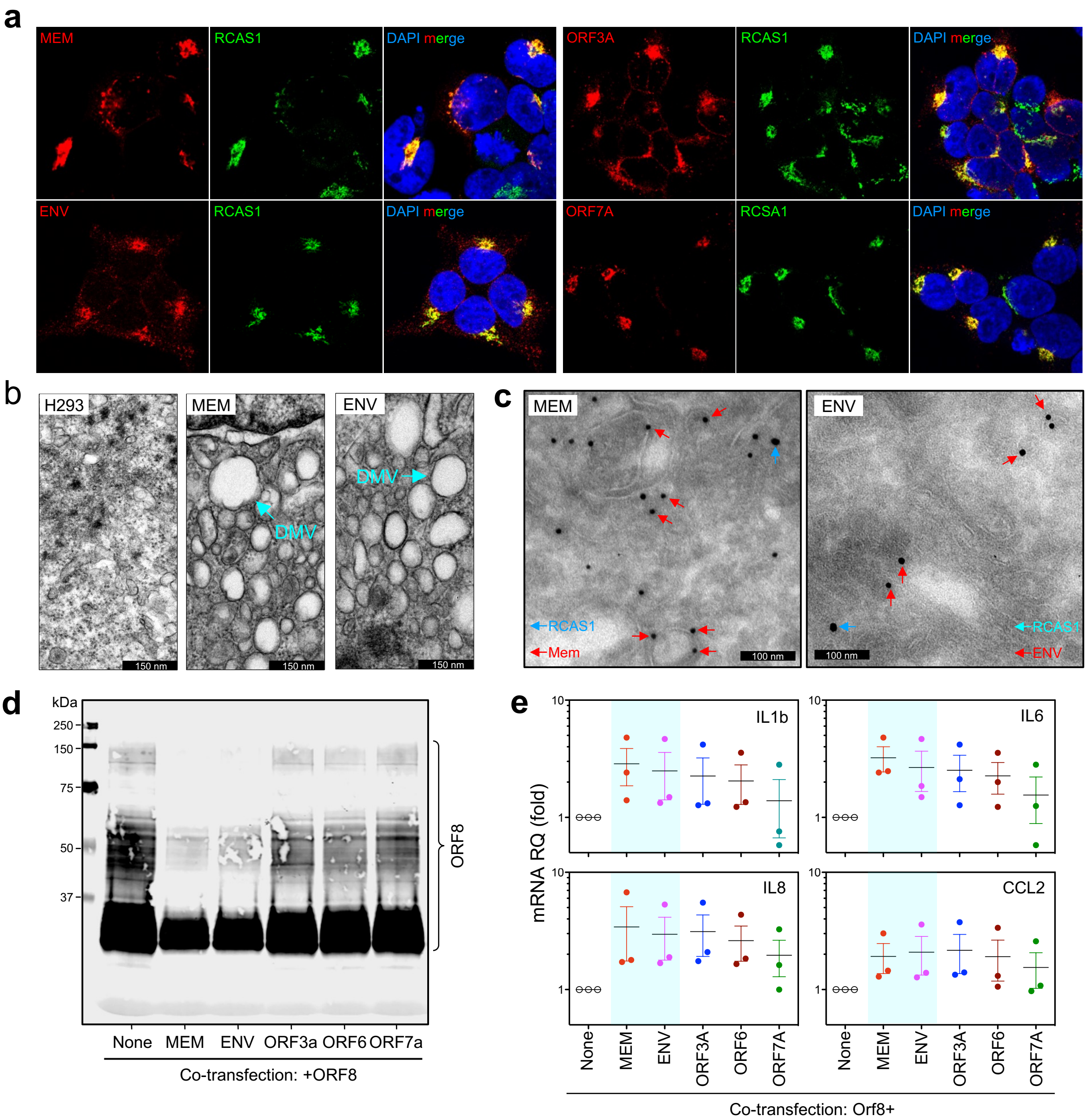


**Supplemental Figure 4.** Membrane targeting accessory proteins of SARS-CoV-2 can modify the structure and function of Golgi leading the neo-formation of double membrane vesicles (DMV) and altered ORF8 glycosylation and cytokine induction activity. **a.b.c.** SARS-CoV-2 membrane targeting proteins MEM, ENV, ORF3A, and ORF7A colocalize with Golgi protein RCAS1 by immunofluorescence staining (**a**), TEM (**b**), and immunogold labeling. **d, e**. Electrophoresis mobility (glycosylation) of ORF8 is affected by co-expressing membrane targeting proteins of SARS-CoV-2 (**d**), leading to the enhancement of their cytokine induction activity (**e**).

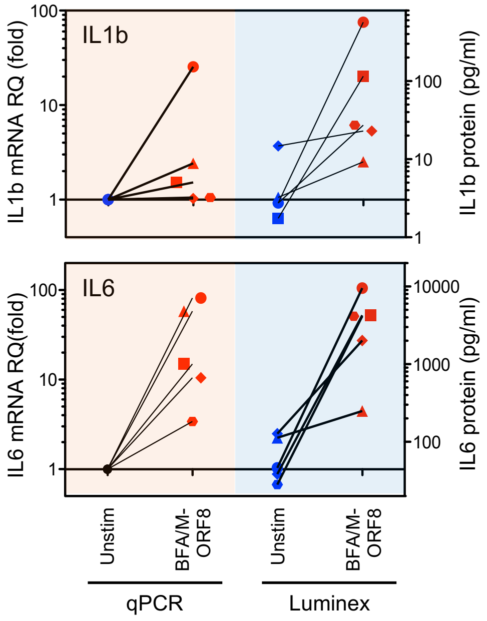


**Supplemental Figure 5. Protein vs mRNA: sensitivity comparison between Luminex methods and qRT-PCR for detecting cytokine expression.** Cytokine IL1b and IL6 proteins in culture supernatants of PBMCs stimulated with CMs from untreated (blue) or BFA/M treated (red) ORF8-expressing HEK293 cells for 3 days were quantitated by Luminex immune assay, while mRNAs of IL1b and IL6 isolated from PBMCs of the same treatment above were quantitated using qRT-PCR.

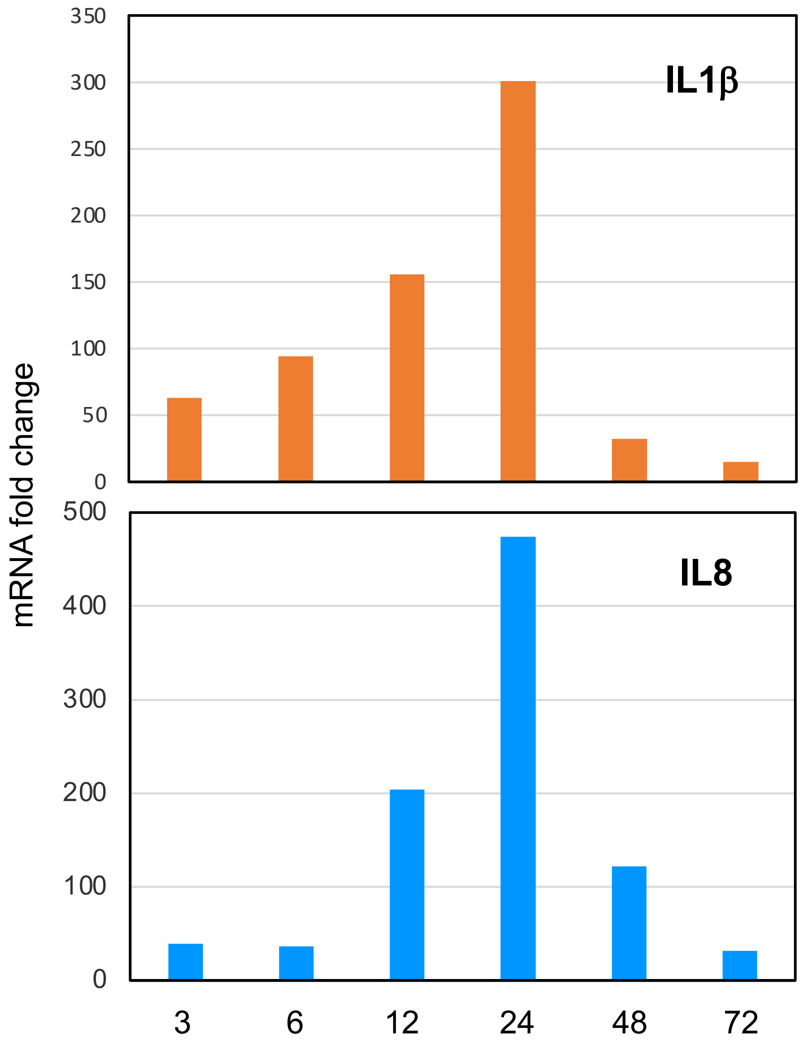


**Supplemental Figure 6.** Kinetics of IL1β and IL8 expression in human monocytes upon stimulation of pure ORF8 at 200 ng/ml. Cells collected at each time point, total RNAs were used for cytokine specific qPCRs, and normalized with HTPR1 gene expression. Data is representative of six samples,

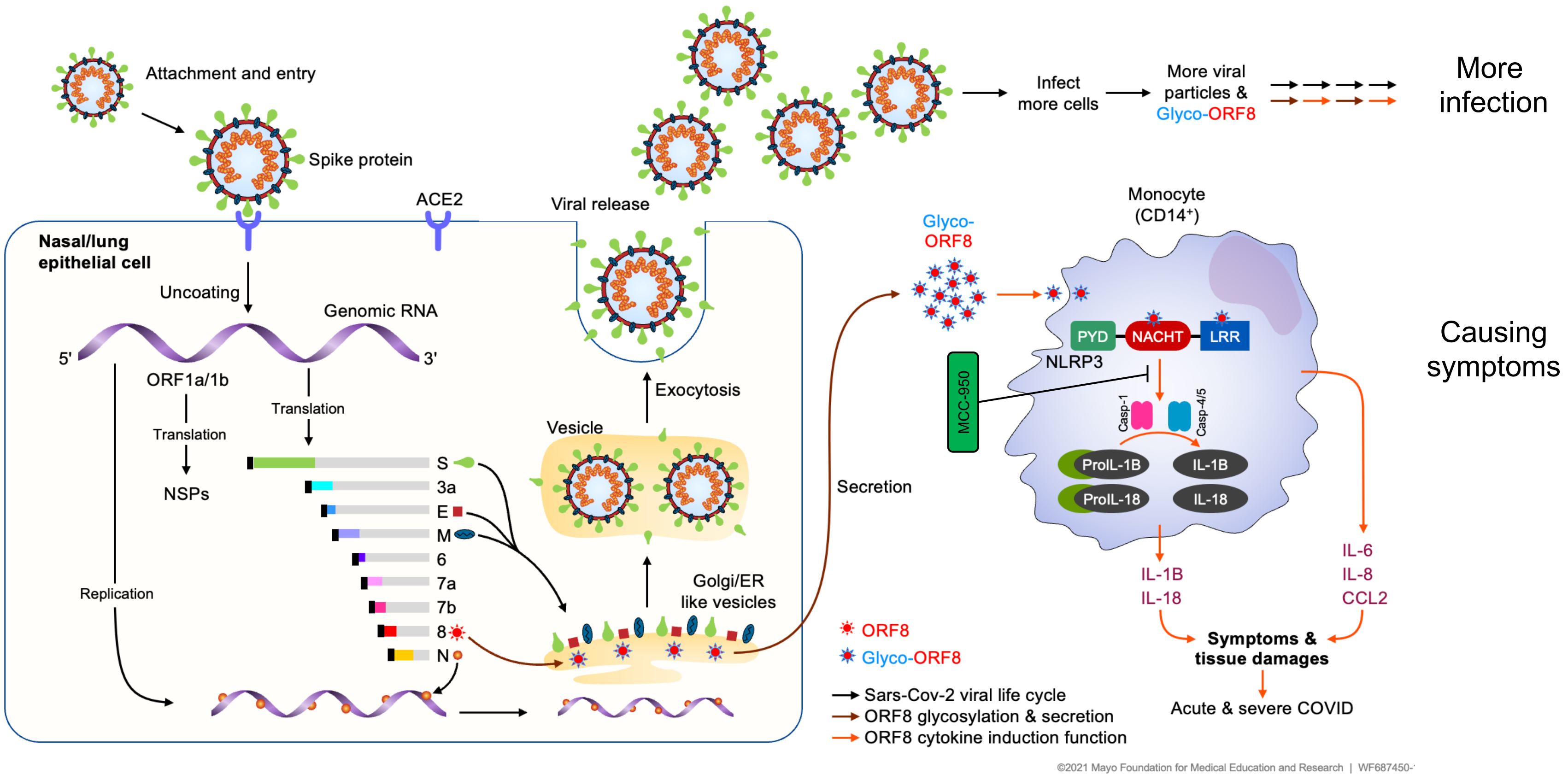


Supplemental Fig. 7. Visual depiction of SARS-CoV-2 life cycle and ORF8 mediated inflammasome response
